# Genomic Characterization and Gene Bank Curation of *Aegilops*: The Wild Relatives of Wheat

**DOI:** 10.1101/2023.07.21.550075

**Authors:** Laxman Adhikari, John Raupp, Shuangye Wu, Dal-Hoe Koo, Bernd Friebe, Jesse Poland

## Abstract

Genetic diversity found in crop wild relatives is critical to preserve and utilize for crop improvement to achieve sustainable food production amid climate change and increased demand. We genetically characterized a large collection of 1,041 *Aegilops* accessions distributed among 23 different species using more than 45K single nucleotide polymorphisms identified by genotyping-by-sequencing (GBS). The Wheat Genetics Resource Center (WGRC) *Aegilops* germplasm collection was curated through the identification of misclassified and redundant accessions. There were 49 misclassified and 28 sets of redundant accessions within the four diploid species. The curated germplasm sets now have improved utility for genetic studies and wheat improvement. We constructed a phylogenetic tree and PCA cluster for all *Aegilops* species together, giving one of the most comprehensive views of *Aegilops*. The *Sitopsis* section and the U genome *Aegilops* clade were further scrutinized with in-depth population analysis. The genetic relatedness among the pair of *Aegilops* species provided strong evidence for the species evolution, speciation and diversification. We inferred genome symbols for two species *Ae*. *neglecta* and *Ae*. *columnaris* based on the sequence read mapping and the presence of segregating loci on the pertinent genomes as well as genetic clustering. The high genetic diversity observed among *Aegilops* species indicated that the genus could play an even greater role in providing the critical need for untapped genetic diversity for future wheat breeding and improvement. To fully characterize these *Aegilops* species, there is an urgent need to generate reference assemblies for these wild wheats, especially for the polyploid *Aegilops*.

**One-sentence summary:** Genotyping *Aegilops* species, the wild relatives of wheat, has revealed high genetic diversity and unique evolutionary relationships among the *Aegilops* and with wheat, giving insight into the effective use of these germplasms for bread wheat improvement.

## Introduction

Global climate change with increasingly variable weather, declining soil quality, and increased biotic and abiotic stresses impede crop production. For instance, an increase in a global mean temperature of a degree Celsius reduces the global wheat yield by 6% (Zhao et al., 2017; Asseng et al., 2015). In this context, the continual genetic improvement of commercial cultivars is needed, including incorporating novel alleles for improved stress tolerance and disease resistance. However, the domestication bottleneck and variety selection practices are major drivers that limit the genetic diversity currently available in the primary gene pool for wheat (*Triticum aestivum* L.) improvement (Haudry et al., 2007). Several past studies have indicated that wild wheat relatives are reliable sources for increasing the genetic diversity in wheat breeding (Lopes et al., 2015).

In this respect, the genus *Aegilops* encompasses the secondary and tertiary gene pool of bread wheat with a central role in wheat evolution and domestication being the donors of B and D subgenomes. As a taxon with high allelic diversity, *Aegilops* species are critically important to provide biotic resistance and abiotic tolerance as well as yield-related genetic loci to wheat (Rakszegi et al., 2020; Kishii, 2019). For instance, *Ae*. *speltoides* harbors agronomically important genes, such as *Sr32* which is effective against the devastating wheat stem rust pathogen Ug99 (Friebe et al., 1996). Similarly, *Ae*. *kotschyi* has been shown to confer leaf and stripe rust resistance with genes *Lr54*, and *Yr37* (Marais et al., 2005), and *Ae*. *biuncialis* possesses a wheat powdery mildew resistance gene (Li et al., 2019). Likewise, the 2NS translocation from *Ae*. *ventricosa* provided multiple disease resistance including root-knot nematode, stripe rust, stem rust, leaf rust and the wheat blast caused by *Magnaporthe oryzae* (Cruz et al., 2016; Gao et al., 2021). Finally, *Ae*. *tauschii* has been frequently used in wheat breeding as the genetic resource for various wheat disease resistance and abiotic-stress tolerance (Suneja et al., 2019).

Though *Aegilops* species hold great potential as genetic resources, limited information is available on the genomic characterization of the genus as a whole. Most of the work to date has been based on cytology, traditional molecular markers, and a limited number of loci, while focusing within a given species. It is complicated by the fact that *Aegilops* species have various ploidy levels and unique genomic compositions and some polyploids have multiple copies of the same sub-genome [e.g. DDM, 6X *Ae*. *crassa*]. Also, reference genomes for only a few *Aegilops* species have been released to date. Therefore, the complicated genomic features and inadequate resources are major challenges for *Aegilops* population studies and more focused, targeted mining of the genetic resources.

These limitations are quickly changing with the recently available genome assemblies of some diploid *Aegilops* such as *Ae*. *tauschii* (Luo et al., 2017), *Ae*. *speltoides* and *Ae*. *longissima* (Avni et al., 2022), *Ae*. *sharonensis* (Yu et al., 2022), *Ae*. *bicornis*, and *Ae*. *searsii* (Li et al., 2022). These genome assemblies are shedding light on *Aegilops* evolutionary and population genetic analysis. Additionally, the high throughput sequencing method such as genotyping-by-sequencing (GBS) (Elshire et al., 2011), which can generate *de novo* genomics variants and is also effective for complex genome species (Poland et al., 2012), has also been proven as an efficient genotyping tool for gene bank collections (Adhikari et al., 2022a).

The Wheat Genetics Resource Center (WGRC) gene bank at Kansas State University has been maintaining myriads of wild wheat accessions under the *Triticum* and *Aegilops* genera. We previously curated the collections of A-genome diploid wheat (Adhikari et al., 2022a) and *Ae*. *tauschii* (Singh et al., 2019a). Thus, the focus of this current study was to characterize the genetic diversity, population structure and genomic composition of the *Aegilops* collection in the WGRC with the curation of the germplasm. Throughout this study, we followed the *Aegilops* species nomenclature by van Slageren (1994) with the exception of *Ae*. *mutica*, and genome symbols were followed as described by Waines and Barnhart (1992). Utilizing variants from genotyping-by-sequencing (GBS), we dissected the genetic and genomic relationships among the 23 *Aegilops* species through phylogenetic clustering, principal component analysis (PCA), population structure analysis, and diversity analysis. We also examined *Aegilops* and wheat genomes relationships through *Aegilops* sequence mapping to the wheat genome and genetic clustering.

## Results

### *Aegilops* Distributions

*Aegilops* species characterized in this study were primarily collected around the fertile crescent, Anatolia, central Asia, northern Africa, and southern Europe (Figure. 1, Supplemental Table S1). Of the five sections, the *Aegilops* section [*Ae*. *umbellulata* (U), *Ae*. *kotschyi* (US), *Ae*. *peregrina* (US), *Ae*. *triuncialis* (CU), *Ae*. *columnaris* (UM), *Ae*. *biuncialis* (UM), *Ae*. *neglecta* (UM, UMN), *Ae*. *geniculata* (MU)] has a much wider distribution from central Asia to northern Africa (Figure. 1). The species of *Cylindropyrum* [*Ae*. *markgraffii* (C), *Ae*. *caudata* (C) and *Ae*. *cylindrica* (CD)] were primarily collected from Uzbekistan, Tajikistan, Kazakhstan, Azerbaijan, and Turkey. The species of the *Comopyrum* [*Ae*. *comosa* (M), *Ae*. *uniaristata* (N)] mainly come from Greece, Turkey and Russia. The *Sitopsis* (S genome) species [*Ae*. *bicornis*, *Ae*. *searsii*, *Ae*. *sharonesis*, *Ae*. *longissima* and *Ae*. *speltoides*] were predominantly collected in Turkey, Israel, Syria, Iraq and Jordan. The *Vertebrata* section species [*Ae*. *tauschii* (D), *Ae*. *crassa* (DM, DDM), *Ae*. *ventricosa* (DN), *Ae*. *juvenalis* (DMU), and *Ae*. *vavilovii* (DMS)] were obtained from central Asia to southern Europe [Figure. 1, Supplemental Table S1]. The *Ae*. *mutica* tested here were from Turkey and Armenia (Supplemental Table S1).

**Figure 1.**
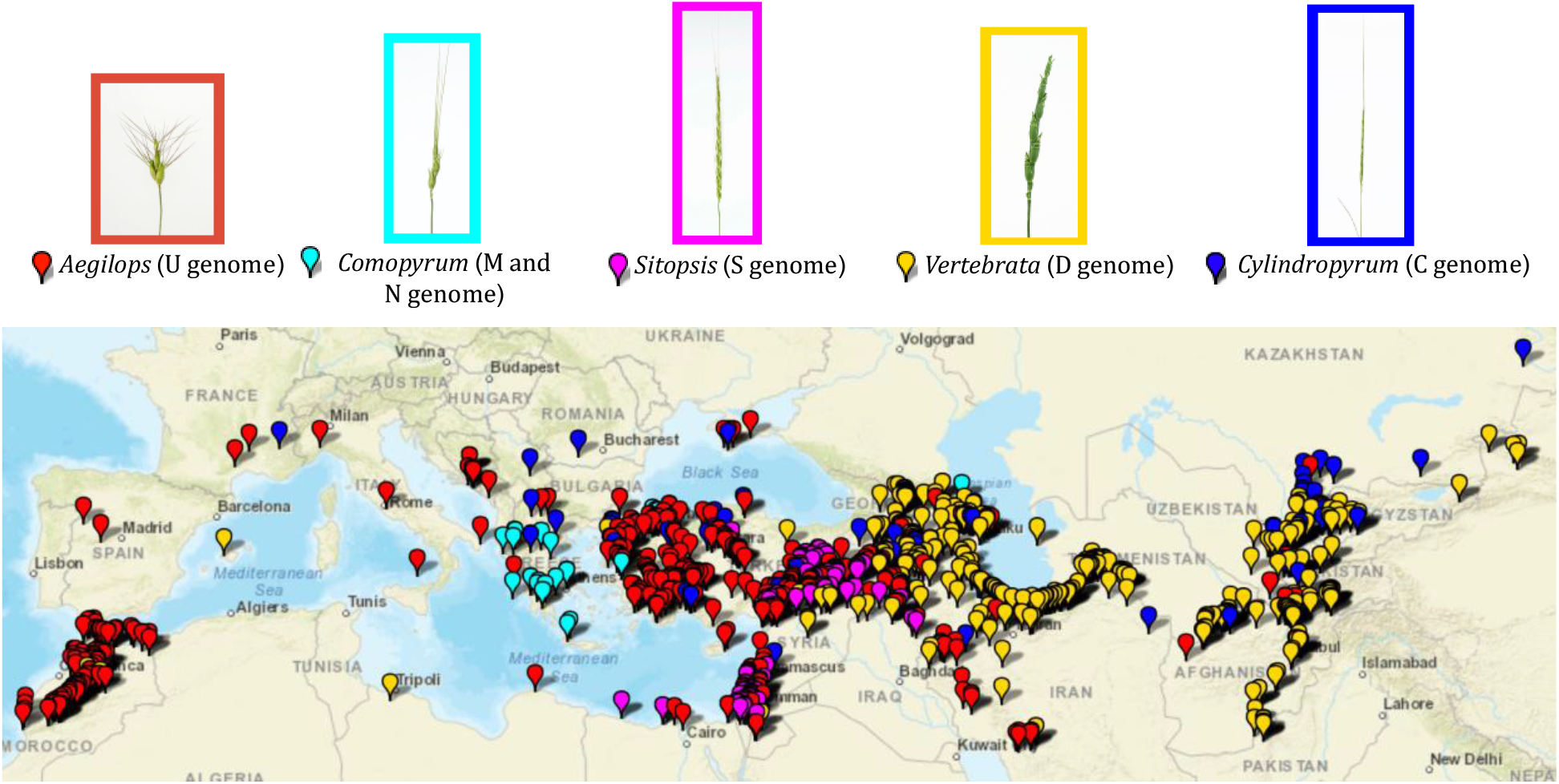
Geographic distribution of the *Aegilops* accessions maintained in the WGRC gene bank. Spike morphologies of representative accessions for the five *Aegilops* sections are shown with the enclosed rectangles. Each section is designated by corresponding color.

### Marker Discovery

We identified 54,667 *de novo* called SNPs for the entire *Aegilops* collections genotyped together. After filtering (MAF > 0.01, missing <30%, heterozygosity <10%) we retained 46879 SNPs (Table 1). We removed 10 accessions (TA2674, TA2633, TA1733, TA11097, TA1740, TA2178, TA2042, TA1739, TA2316, TA2296) with high rate of missing call (> 80%). When we separated the genotyping information per species, we identified filtered segregating SNPs in the range of 1,483 for *Ae*. *searsii* to 14,322 for *Ae*. *speltoides* (Table 1). We also generated other SNP genotyping matrices for analysis-specific purposes, such as for particular species’ genetic relations and for genetically identical accession determination (Supplemental Table S3).

**Table 1.** *Aegilops* species with number of accessions, number of segregating loci and the Nei’s diversity indices. (*) The *Ae*. *mutica* being cross-pollinated we used many different samples from a single accession (s), so total of 54 plants rather than accessions.

| Species | # Accessions | Segregating loci | Nei's index |
| --- | --- | --- | --- |
| All Collection | 1041 | 54668 | 0.104 |
| *Ae. tauschii* | 47 | 3369 | 0.024 |
| *Ae. vavilovii* | 6 | 9955 | 0.093 |
| *Ae. mutica* | 54* | 8094 | 0.053 |
| *Ae. ventricosa* | 17 | 5828 | 0.05 |
| *Ae. uniaristata* | 24 | 5416 | 0.019 |
| *Ae. umbellulata* | 58 | 3391 | 0.015 |
| *Ae. triuncialis* | 199 | 8601 | 0.032 |
| *Ae. speltoides* | 97 | 14322 | 0.072 |
| *Ae. sharonensis* | 9 | 2224 | 0.019 |
| *Ae. searsii* | 18 | 1483 | 0.013 |
| *Ae. peregrina* | 33 | 7981 | 0.053 |
| *Ae. neglecta* | 71 | 11931 | 0.062 |
| *Ae. markgrafii* | 16 | 3474 | 0.022 |
| *Ae. longissima* | 14 | 3043 | 0.023 |
| *Ae. kotschyi* | 24 | 6876 | 0.053 |
| *Ae. juvenalis* | 9 | 8796 | 0.081 |
| *Ae. geniculata* | 143 | 8248 | 0.038 |
| *Ae. cylindrica* | 79 | 6173 | 0.046 |
| *Ae. crassa* | 32 | 8999 | 0.074 |
| *Ae. comosa* | 17 | 3388 | 0.025 |
| *Ae. columnaris* | 12 | 5382 | 0.041 |
| *Ae. biuncialis* | 52 | 7819 | 0.042 |
| *Ae. bicornis* | 13 | 1493 | 0.012 |

### Gene Bank Curation Misclassified accessions

The phylogenetic clustering and PCA enabled us to identify and correct the classification of 49 accessions (Figure 2 and Supplemental Table S2). Most of the misclassified accessions were observed within tetraploid *Aegilops*. Twelve accessions that were previously considered as *Ae*. *triuncialis* were now identified as different *Aegilops*, whereas nine accessions that were classified as different *Aegilops* species are now re-identified as *Ae*. *triuncialis* (Supplemental Table S2). Similarly, 11 accessions identified as *Ae*. *neglecta* were now genetically identified as different *Aegilops*. The other misclassified example includes four accessions of each of *Ae*. *geniculata* and *Ae*. *vavilovii* (Supplemental Table S2). A few misclassified accessions of diploid *Aegilops* included *Ae*. *umbellulata* (2), *Ae*. *markgrafii* (2) and *Ae*. *searsii* (1) (Figure 2). The classes of all misclassified accessions were updated prior to the downstream population genomic analysis.

**Figure 2.**
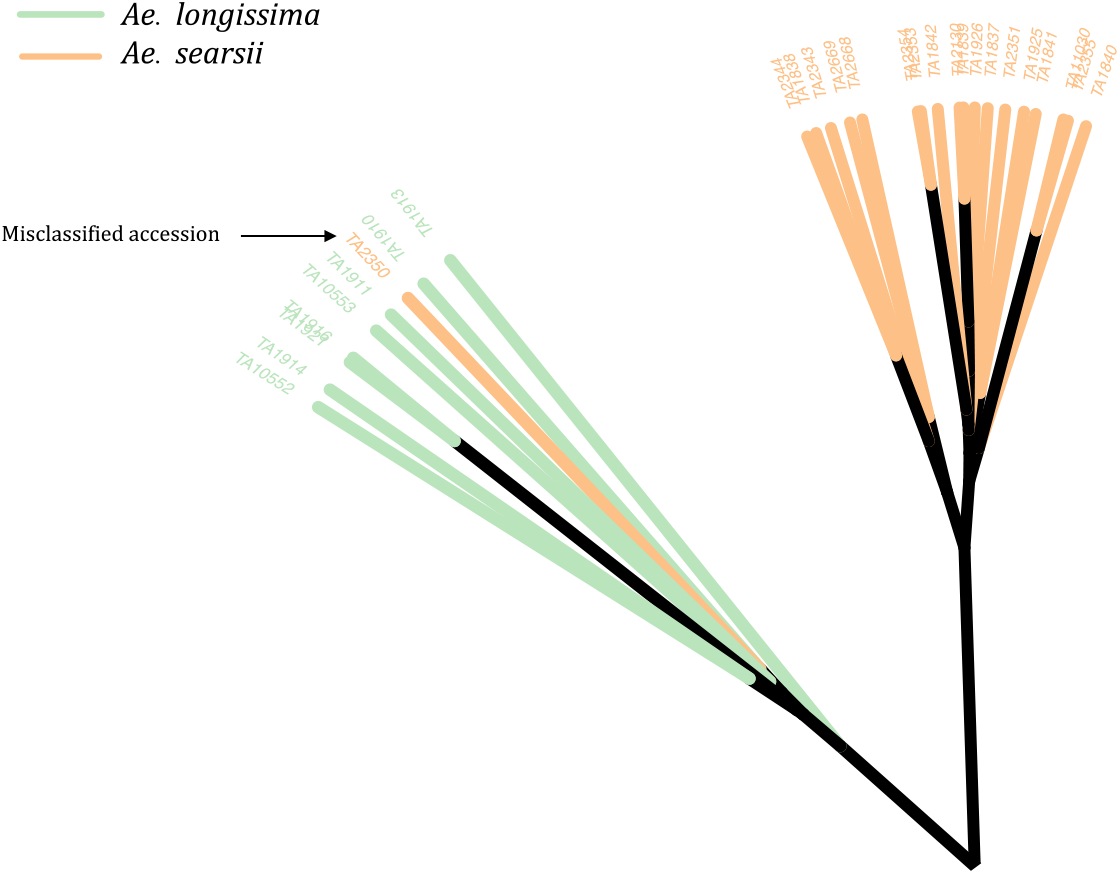
An unrooted neighbor-joining (NJ) tree with an example of a misclassified accession (TA2350) in the WGRC gene bank. The genetically clustered clades were colored based on the morphological classes of the accessions and visually accessed. The misclassified accession TA2350, which was previously grouped under *Ae*. *searsii* (orange clade) was re-classified as *Ae*. *longissima* (green).

### Genetically Identical Accessions

The gene bank curation discovered total 28 genetically identical accessions in *Ae*. *tauschii* and four members of the *Sitopsis* section (Supplemental Table S2). There were no pairs of *Ae*. *speltoides* accessions that have allele matching above 95%. Of 28 duplicated accessions, 17 were from *Ae*. *tauschii*, even though we only had a total of 47 *Ae*. *tauschii* accession for this experiment (Supplemental Table S2). In our previous study, we also reported many genetically identical accessions in *Ae*. *tauschii* collection (Singh et al., 2019a).

### Phylogenetic Clustering, PCA, and Population Structure

The unrooted neighbor-joining (NJ) phylogenetic tree with all tested *Aegilops* accessions gave clear separation of species as the branches of clades and sub-clades differentiated all 23 species and the relevant groups (Figure 3). We observed the species sharing genomes as closely related clades, such as *Ae*. *kotschyi* and *Ae*. *peregrina* (SU), and *Ae*. *geniculata* and *Ae*. *biuncialis* (UM) clustered into respective primary clades (Figure 3). Overall, there were three primary clades; i) the first clade consisted of *Ae*. *speltoides* and *Ae*. *mutica*; ii) the second clade has four diploids of *Sitopsis* (except *Ae*. *speltoides*), *Ae*. *tauschii*, and D genome polyploids (except *Ae*. *cylindrica*); iii) the third primary clade has all other species, including M, N, C and U genome diploids and polyploids (Figure 3).

**Figure 3.**
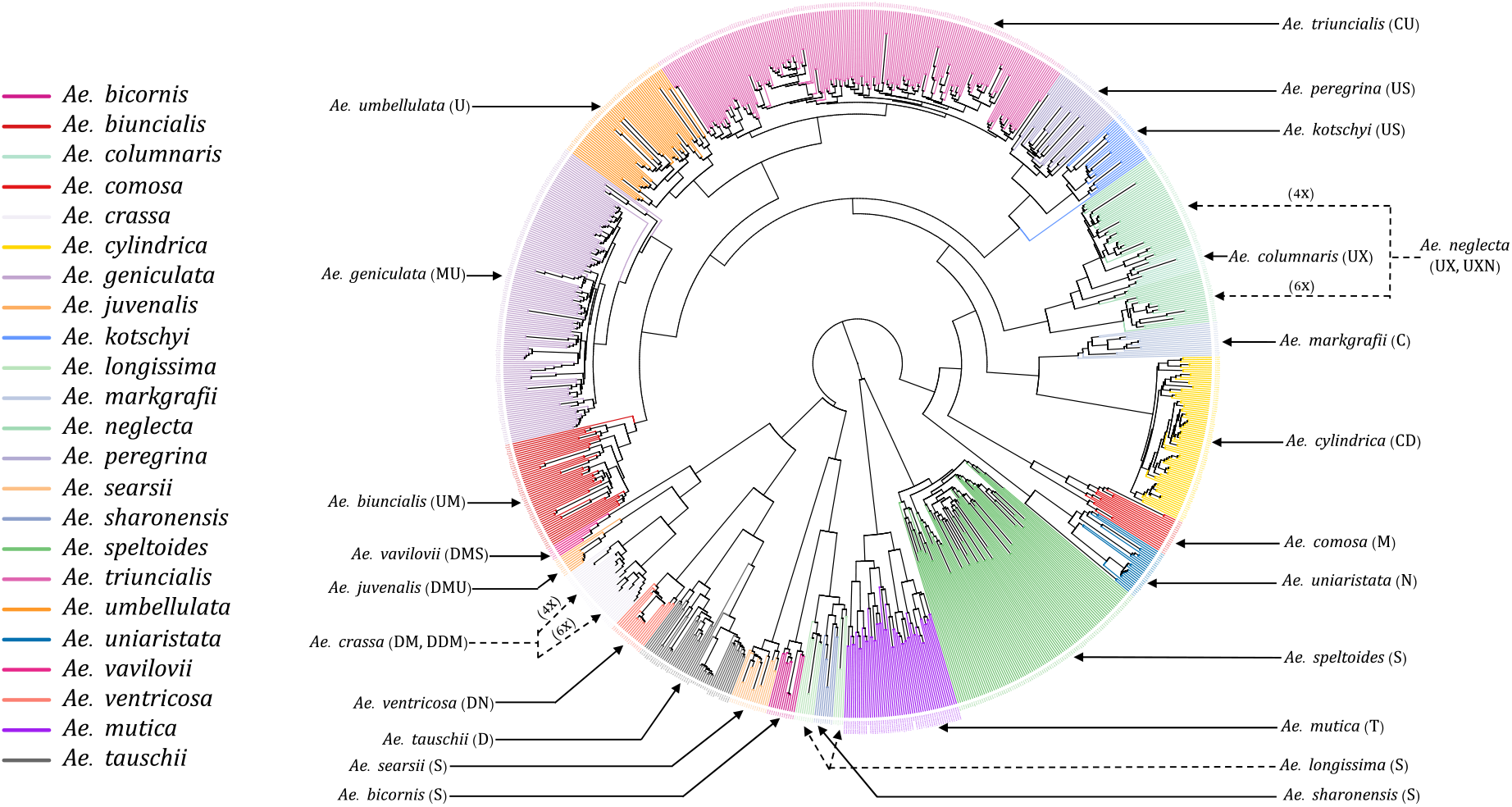
An unrooted neighbor-joining (NJ) tree of 23 different *Aegilops* species. The tree branches were colored based on the accessions genetic grouping after adjusting the misclassified accessions classes. The genome signs of each of the species were annotated along with their names as indicated by solid and dotted arrowheads.

The hexaploid (6X) and tetraploid (4X) species within a clade, such as *Ae*. *neglecta* and *Ae*. *crassa* were grouped separately by ploidy. The ploidy levels of these genetically clustered sub-groups (6X and 4X) were also verified using chromosome counting (Figure S1) following Koo et al. (2017). The chromosome numbers of some accessions of *Ae*. *crassa* (Figure S5) were also confirmed with the published data (Badaeva et al., 1998).

Principal component analysis (PCA) also grouped the *Aegilops* species commensurate with the phylogenetic analysis. The first and second principal components (PC1 and PC2) explained about 17% and 14% of the variations among the *Aegilops*, respectively. PC1 separated *Ae*. *speltoides* from other polyploids and diploids (Figure 4), while the PC2 primarily differentiated *Ae*. *tauschii* and *Ae*. *speltoides*, the D genome donor to wheat and the potential sister group of the wheat B genome donor, respectively. As in phylogenetic analysis, PCA grouping also divided the 4X and 6X accessions of the *Ae*. *neglecta* and *Ae*. *crassa* (Figure 4).

**Figure 4.**
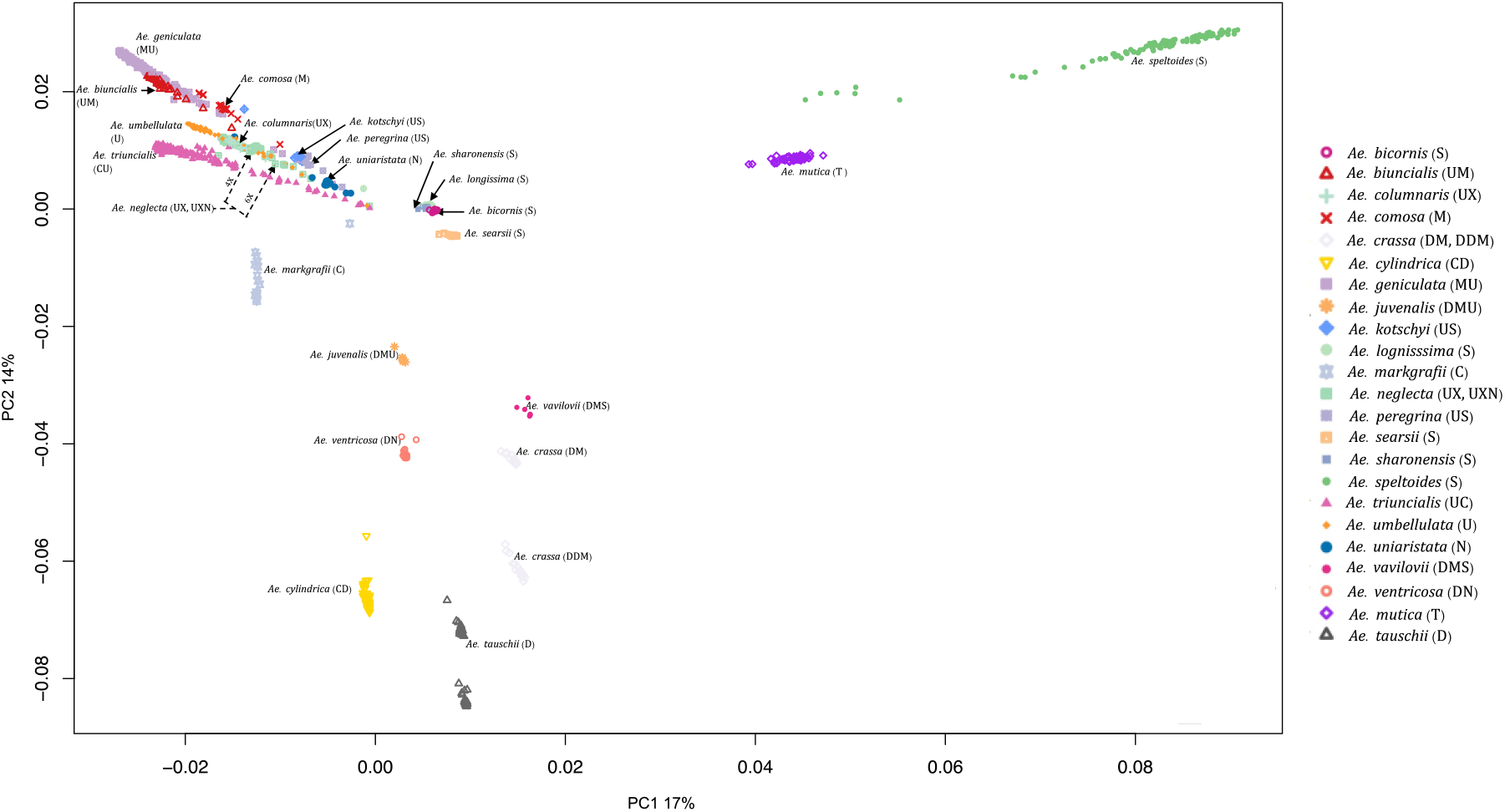
Principal component analysis (PCA) plot for all 23 *Aegilops* species with the first two principal components (PCs). The 23 *Aegilops* species were grouped and colored based on their species and genome compositions.

### Population Genomics of *Sitopsis* and *Ae*. *mutica*

As we observed the separation of four *Sitopsis* members with *Ae*. *speltoides* and *Ae*. *mutica*, we separately examined the population of these species using reference-based variants from the *Ae*. *speltoides* genome assembly. The constructed phylogenetic tree distinctly divided the S-genome diploids into two large clades, one representing *Ae*. *speltoides* and the other with the remaining four *Sitopsis* (Figure 5). The genetic clustering corresponded to the historical sub-section division of the section as *Truncata* (*Ae*. *speltoides*) and the *Emarginata*. We also observed that the *Ae*. *mutica* (T genome) clustered closer to *Ae. speltoides* both in PCA and phylogenetic analysis (Figure 5). The relationships among *Sitopsis* group and *Ae*. *mutica* were further verified by computing pairwise Nei’s F_ST_ (Nei, 1987), where we observed *Ae*. *mutica* has the closest genetic relationship [lowest F_ST_ (0.65)] with *Ae*. *speltoides*, closer than any other members of the *Sitopsis* (Supplemental Table S4). Hence, all these analyses support that *Ae*. *mutica* as the sister taxon to *Ae*. *speltoides* and it is an *Aegilops* species.

**Figure 5.**
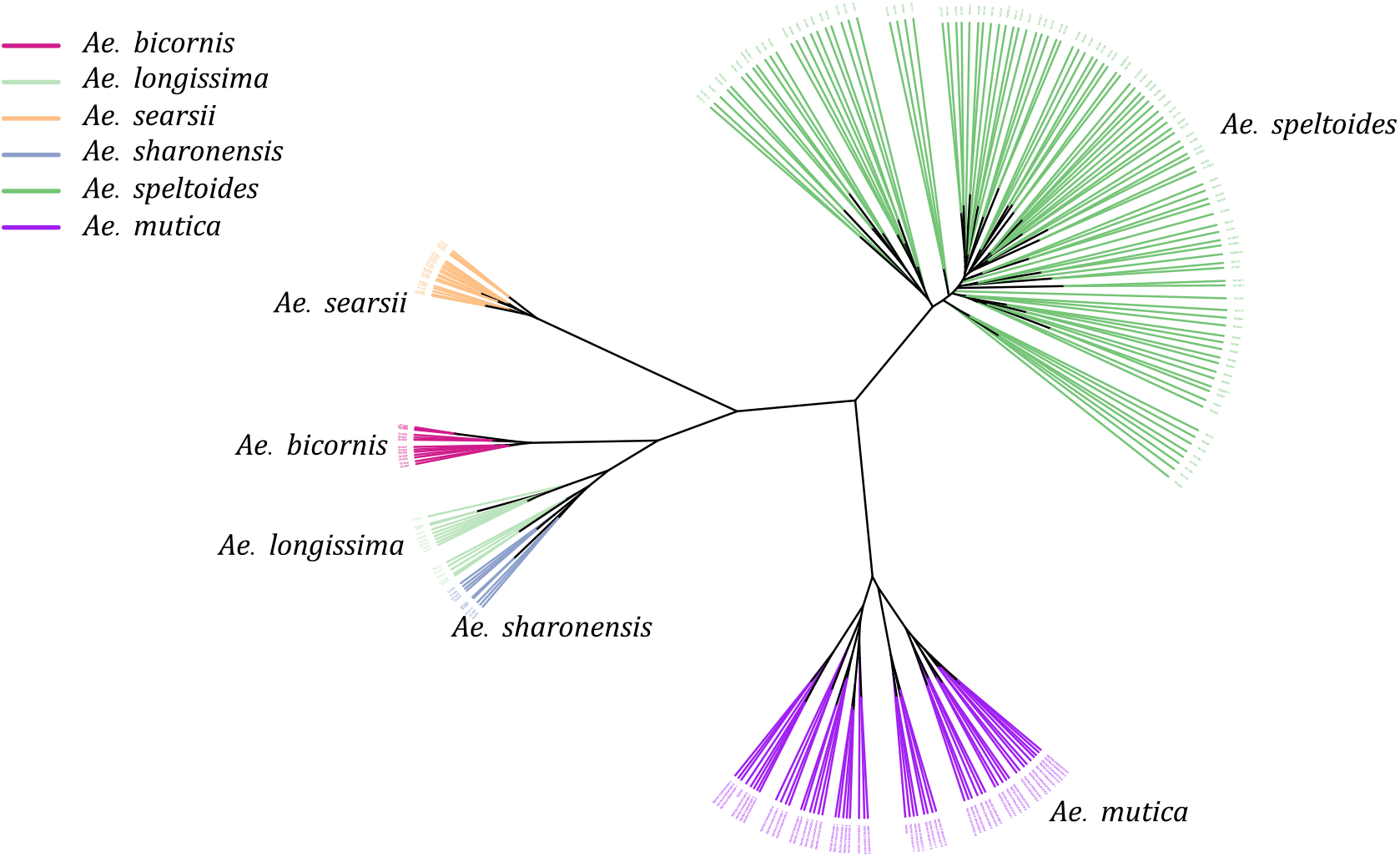
An unrooted Neighbor-Joining tree of five *Aegilops* species including *Sitopsis* section members (S genome) and *Ae*. *mutica* (T genome).

Further, within the S-genome diploids, the *Ae*. *speltoides* and *Ae*. *searsii* had the most genetic differentiation with the highest F_ST_ value 0.88 (Supplemental Table S4). However, the pairwise F_ST_ indicated that *speltoides* is genetically almost equally and highly differentiated from all other S-genome diploids (*Emarginata*) (Supplemental Table S4).

Population structure analysis of S-genome diploids matched with the phylogenetic tree and pairwise F_ST_ analysis. At K=2, there was a differentiation between *Ae*. *speltoides* and the rest of the *Sitopsis*, while at K=3, *Ae*. *searsii* also differentiated from the rest of the *Sitopsis* (Figure 6). At K=7, *Ae*. *bicornis* accessions separated from others and then no new differentiation was observed until K=12. Both in the phylogenetic tree and in population structure analysis, the *Ae*. *longissima* and *Ae*. *sharonensis* appeared as highly genetically similar groups (Figure 5 and 6). In fact, there were no population differentiation between these two species at any level of K. The pairwise F_ST_ values also confirmed that these two species have the lowest pairwise F_ST_ = 0. 006 (Supplemental Table S4) and the population differentiation is very low. Further, two sub-groups within *Ae*. *speltoides*, *var*. *speltoides* and *var*. *ligustica*, also did not differentiate at any levels of K in the population structure analysis (Figure 6) and the PCA (Figure S2). However, within *Ae*. *speltoides* a few admixtures were observed and were differentiated for their geographical origins (Figure 6).

**Figure 6.**
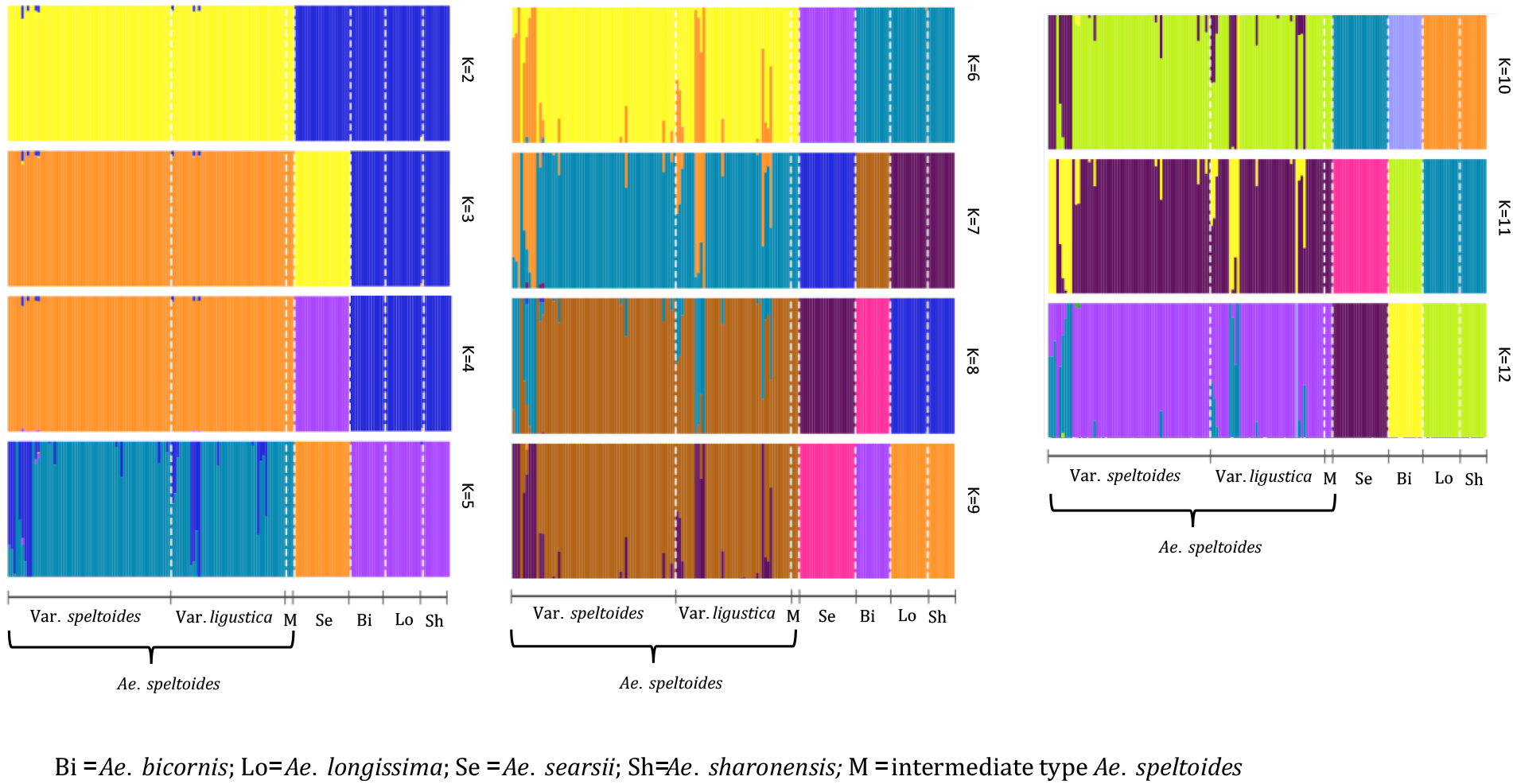
The Population structure of S-genome diploids *Aegilops*, where the value of K and colors of the bars indicate the description of the groups. Each color represents a population and each bar with more than one color indicates the admixtures with the admixture proportions as represented by the proportion of each color.

### *Ae*. *umbellulata* and U-genome Tetraploids

Most of the tetraploid *Aegilops* have the U genome, therefore, understanding the genetic relationship among members of the U-genome clade gives insight into a large set of taxa in the genus. Phylogenetic clustering of these species only showed two larger clades, where one was represented by *Ae*. *triuncialis* (UC) and the other had all remaining tetraploids (Figure 7). The diploid *Ae*. *umbellulata* sits on the intermediate position between the larger clades. Although the variants were only called on U-genome (*Ae*. *umbellulata*) *de novo* reference, the tetraploids distinctly grouped for their genomic compositions. The tetraploid species *Ae*. *pregerina* and *Ae*. *kotschyi* (US genome), *Ae*. *neglecta* and *Ae*. *columnaris* (traditionally assigned as UM), and the UM genome tetraploids *Ae*. *biuncialis* and *Ae*. *geniculata* formed a separate clade and sub-clades (Figure 7). Also, we observed the splitting of *Ae*. *umbellulata* accessions into smaller clades. With a few exceptions as noted below, these phylogenies largely agree with previous genome designations.

**Figure 7.**
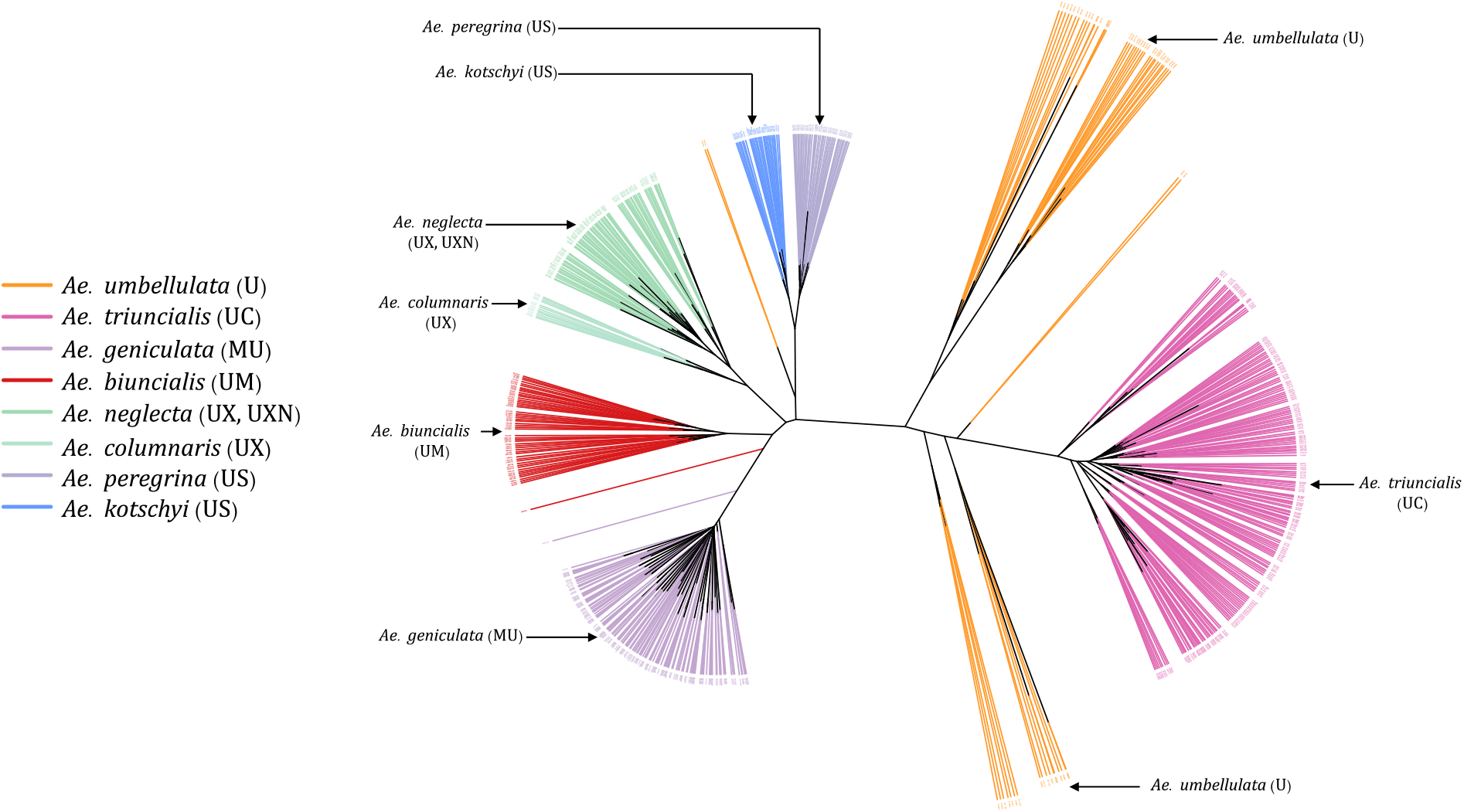
An unrooted neighbor-joining (NJ) tree for *Ae*. *umbellulata* and U genome containing tetraploids within the genus *Aegilops*.

### Genome Symbols of *Ae*. *columnaris* and *Ae*. *neglecta*

*Ae*. *columnaris* and *Ae*. *neglecta* formed a different clade than the other tetraploids with U and M genomes such as *Ae*. *geniculata* (UM) and *Ae*. *biuncialis* (MU) in both phylogenetic clustering and PCA (Figures 3, 4, 7 and Supplemental Figure S3). The comparative positions of these tetraploids with other tetraploids in the genetic cluster indicated that these two tetraploids must be given unique genome symbols than the *Ae*. *geniculata* and *Ae*. *biuncialis* (Supplemental Figure S3). Thus, we hypothesized that *Ae*. *columnaris* and *Ae*. *neglecta* do not carry the M genome. The absence of M genome in *Ae. columnaris* and *Ae. neglecta* accessions was further confirmed by computing total reads mapped and total variants called on M-genome (*Ae*. *comosa* mock reference) and U genome (*Ae*. *umbellulata* mock reference) (Supplemental Figure S4 and Supplemental Table S5). All four tetraploid species including *Ae. columnaris* and *Ae. neglecta* along with *Ae. geniculata* and *Ae. biuncialis* exhibited an equal percentage of overall reads alignment (∼ 38%) on the U genome, whereas the percentage read alignment of *Ae. columnaris* and *Ae. neglecta* on M genome was low (∼21%) as compared to the alignment of *Ae. geniculata* and *Ae. biuncialis* reads (∼ 38%). We also noticed that a few *Ae*. *comosa* segregating loci were mapped for *Ae*. *columnaris* (10%) and *Ae*. *neglecta* (24%) on the M genome. In contrast, *Ae*. *biuncialis* had 50% and *Ae*. *geniculata* had 46% M-genome loci. Hence, the proportion of mapped reads and loci also suggested the *Ae. neglecta* and *Ae*. *columnaris* must have the U genome, but a different second sub-genome than M. Thus, we proposed that *Ae. columnaris* and *Ae. neglecta* genome formulas are most likely UX (X, the unknown genome) or UXN in hexaploid form as proposed based on cytology (Badaeva et al., 2018; Dvorak, 1998).

### *Aegilops* Species Diversity

For the whole collection we obtained 54,667 SNPs, which were skewed to low minor allele frequency (MAF) as expected for a diverse population such as this (Supplemental Figure S6). Despite the differences in population size, the total segregating loci for the species or groups were mostly dependent on the ploidy levels and the reproductive biology (inbreed vs outcrossing) (Table 1). The polyploids and out-crossing species had a higher number of segregating loci as compared to other diploids (Table 1). The MAF of most of the loci of the partially cross-pollinated species such as *Ae*. *speltoides* had a higher frequency (Supplemental Figure S7) than the MAF of the loci for the entire *Aegilops* collection (Supplemental Figure S6).

The Nei’s diversity indices also followed the pattern of segregating loci which were greater in polyploid and cross-pollinated species. We computed Nei’s diversity index for the entire collection as 0.10 (Table 1). Of all 23 species, *Ae*. *bicornis* had the lowest Nei’s diversity index (0.012) followed by *Ae*. *searsii* (0.013), and *Ae*. *umbellulata* (0.015). Among the diploids, the *Ae*. *speltoides* had the highest Nei’s diversity (0.072) which was followed by *Ae*. *mutica* (0.053). Among the tetraploids, the *Ae*. *triuncialis* had the lowest diversity index (0.032) while the *Ae*. *neglecta* had the highest diversity index (0.062). The hexaploid species *Ae*. *vavilovii* has the highest Nei’s diversity index value among all 23 species analyzed in the experiment (Table 1). This increased diversity can be attributed to various factors such as multiple gene copies, hybridization during speciation, increased mutation rates, and more opportunities for recombination due to the presence of multiple genomes.

### Wheat and *Aegilops* Genomes

The genetic clustering between wheat and all diploid *Aegilops* showed that *Ae*. *tauschii* is the nearest extant *Aegilops* to the bread wheat (Supplemental Figure S8). The genetic cluster clearly showed that *Ae*. *speltoides* is not closer to wheat as *Ae*. *tauschii* and even other diploids, and supporting that *Ae*. *speltoides* is likely not the direct progenitor of the wheat subgenome B (Supplemental Figure S8).

However, the *Ae*. *speltoides* read depth mapping and SNPs detection occurred at its maximal on the wheat subgenome B (Figure 8), indicating the species as the sister group of wheat B genome progenitor. Further, the other members of the *Sitopsis* group clustered between *Ae*. *speltoides* clade and the clade with *Ae*. *tauschii* and the wheat subclades in the phylogenetic tree (Figure S8). Consistent with the genetic clustering, their maximum reads mapping and SNP detection also occurred at subgenome D and B chromosomes (Supplemental Figures S8 and S9), suggesting the four members of *Sitopsis*, except *Ae*. *speltoides*, have very strong genomic relationships with both D and B subgenomes.

**Figure 8.**
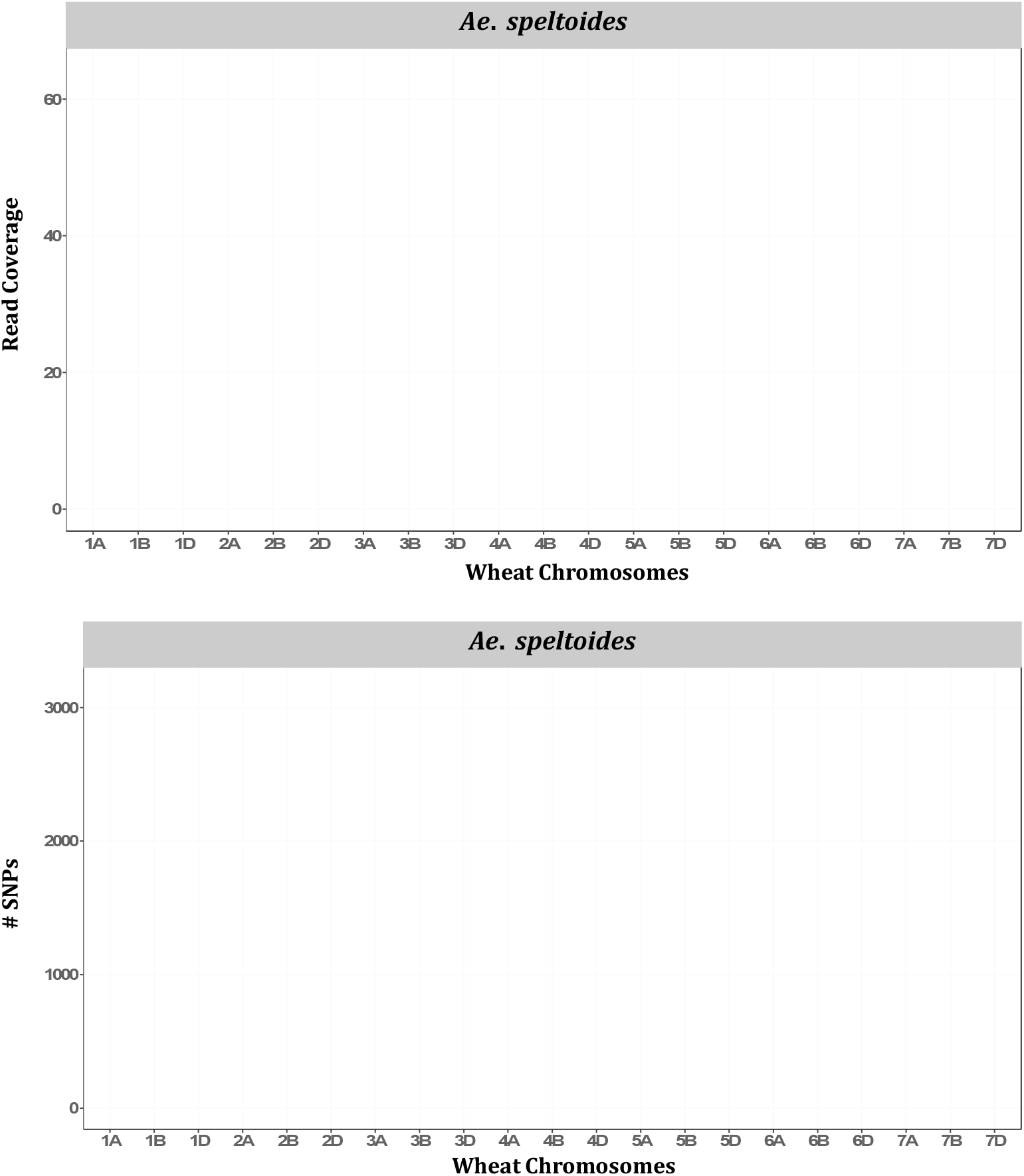
Bar charts showing genomic relations between *Ae*. *speltoides* and wheat. The average number of *Ae*. *speltoides* sequence reads mapped per Mb of the wheat genome (upper panel), and numbers of *Ae*. *speltoides* variants mapped on the respective wheat chromosomes (lower panel).

Similarly, the U genome diploid (*Ae*. *umbellulata*) highest proportion of sequence reads were mapped on wheat chromosomes of the D subgenome followed by A and B (Supplemental Figure S11). Exceptionally, a slightly higher proportion of reads were mapped on 2A than the 2D. The pattern of SNP detection was exactly the same as read mapping, indicating that wheat subgenome D is the closest to the U genome of the *Aegilops*. However, relations between the wheat A genome and the *Aegilops* U genome can’t be overlooked as reasonably higher reads and loci were mapped on the A genome as compared to the wheat B genome (Supplemental Figure S11). Likewise, the highest number of reads and SNPs were mapped on wheat subgenome D for the N genome diploid (*Ae*. *uniaristata*) (Supplemental Figure S12), for the M genome diploid (*Ae*. *comosa*) (Supplemental Figure S13) and C genome diploid (*Ae*. *markgraffii*) (Supplemental Figure S14). These observations suggested that the N, M and C genomes of *Aegilops* were also genetically closer to the D subgenome than A and B.

Interestingly, the *Ae*. *mutica* accessions when mapped to the wheat subgenomes showed higher sequence read and loci mapped on the wheat D subgenome (Supplemental Figure S15). The read and loci mapping pattern was unchanged even we replaced wheat D subgenome chromosomes with *Ae*. *tauschii* chromosomes. Nevertheless, all types of population grouping within *Aegilops* (Figures 3-5 and Supplemental Figure S8) evidently showed that *Ae*. *mutica* is a sister group of *Ae*. *speltoides* and still a member of B lineage. Some recent studies based on whole genome sequencing data have also reported a higher sequence read and loci mapping of *Ae*. *mutica* on the wheat D subgenome than others (Grewal et al., 2022; Li et al., 2022).

## Discussions

### Multi-species Diverse *Aegilops* Collection and Gene Bank Curation

In this study, we genotyped over a thousand accessions representing almost all species of the genus *Aegilops* from the full range of their natural distributions under the van Slageren (1994) nomenclature, with missing only *Ae*. *caudata*. We curated the WGRC gene bank *Aegilops* collection, giving curated germplasm sets that are ready to screen for the novel alleles and utilize in the breeding program. The misclassified accession were confirmed with multiple analyses including phylogenetic clustering of the whole population, species or genome-specific populations and PCA, therefore there is strong support for the genotype-based identification of these misclassified accessions (Supplemental Table S1). Since the genotype-based clustering evidently differentiated the hexaploid and tetraploid accessions within the species like *Ae*. *crassa* and *Ae*. *neglecta*, we can also provide the ploidy levels information as a means of within-species classification and update the gene bank database.

Here we identified the redundant accessions in the species with variants called directly on reference genome assemblies. This gives increased power and accuracy in variant calling. Therefore, we suggest the re-assessment of genetically redundant accessions for other *Aegilops* species in the future when reference assemblies are available. For the polyploid *Aegilops,* reference variant calling can be done whenever the component species reference genomes are available using a combined reference genome or independent variant calling to each genome. Although we do not suggest discarding the duplicated accessions identified here, we strongly suggest for considering these results when utilizing the collection, such as screening the accessions for disease resistance or developing introgression populations. Overall, the gene bank curation helps in management, preservation and utilization of the germplasms (Volk et al., 2021; Singh et al., 2019a).

### *Aegilops* Population Analysis

This is the most comprehensive *Aegilops* population genetic study reported so far with over 45 thousand *de novo* filtered SNPs and reference-based variants. In the study, we took advantage of recently completed chromosome-scale genome assemblies of diploid *Aegilops* (Li et al., 2022; Avni et al., 2022; Yu et al., 2022; Wang et al., 2021). Until now, the lack of genomic resources including reference assemblies has been a major issue hindering the species population genomic analysis. Therefore, future genomic studies on *Aegilops* must focus on generating more genomic resources for other diploids and polyploids. With a larger population and thousands of genomic variants, the population grouping we observed here was at the finest level, enabling us to differentiate the 4X and 6X accessions within a species (Supplemental Figure S1).

### Ae. speltoides, other Sitopsis and Ae. mutica

Our genetic analysis supports that the *Ae*. *mutica* requires no genus-level separation from other *Aegilops* as van Slageren (1994) suggested. It is genetically an *Aegilops* taxon closer to *Ae*. *speltoides* (Figures 4 and 5). This is in agreement with recent reports (Li et al., 2022; Bernhardt et al., 2020). Further genomic analysis may require high coverage genomic data and a greater number of samples to better understand the relationship among *Ae*. *mutica* and other diploid *Aegilops*. Additionally, the genetic differences we observed here between the *Truncata* (*Ae*. *speltoides*) and *Emarginata* (four other) *Sitopsis* were greater, therefore, the redefinition of the section *Sitopsis* could be desirable. One of the ideas could be the separation of *Ae*. *speltoides* from the rest of the four *Sitopsis* members and regrouping the *Ae*. *speltoides* with *Ae*. *mutica* (Figures 3-5 and Supplemental Figure S8).

We also showed that the *Ae*. *sharonensis* and *Ae*. *longissima* have very high genetic similarities or a low genetic differentiation (F_ST_ = 0.006) and are most likely the sub-species of the same species. Also, both of these species are equally distant from *Ae*. *speltoides*. The finding is also supported by the latest study, where Avni et al. (2022) reported that the genomes of these two species are highly similar with identical genome sizes and also share 292 orthogroups.

In this study, we observed a little genetic difference between the two sub-taxa of *Ae*. *speltodies*; var. *speltoides* and *ligustica* with no population differentiation (Figure 6 and Supplemental Figure S2), in accordance with several past studies. These two subgroups of *speltoides* have distinct spike morphology and mode of seed dispersal, but also exhibit similar karyotype structure, producing fully fertile hybrid and mixed stands of two types naturally exhibits (Zohary and Imber, 1963). A single locus *Lig* on chromosome 3S governs the spike morphology of these two sub-groups (Luo et al., 2005); otherwise, they are highly genetically similar.

### U-Genome species, some tetraploid genome Symbols and polyploid *Aegilops*

The U genome tetraploids and its progenitor *Ae*. *umbellulata* genetic clustering revealed the unique relationships among the species. We observed the *Ae*. *umbellulata* accessions split into sub-groups in such a way that some accessions were clustered closer to *Ae*. *triuncialis* clade whereas some other accessions reposed near the other tetraploid clades (Figure 7), suggesting the potential unique *Ae*. *umbellulata* ancestries for the two groups.

In this study, we found further evidence that the *Ae*. *columnaris* and *Ae*. *neglecta* genome symbols should not contain M genome (Supplemental Figures S3 and S4, and Supplemental Table S5) with sequence read and loci mapping data and the phylogenetic clustering (Supplemental Figure S3). Based on the cytology-based approaches (Badaeva et al., 2018; Badaeva et al., 2004; Dvorak, 1998; Resta et al., 1996), studies have already discussed this genome symbols issue and suggested the symbol ‘X’ (Resta et al., 1996). Several pieces of evidence, such as a low chromosome pairing in hybrids of *Ae*. *columnaris* x *Ae*. *comosa* (M genome progenitor), variation of the repetitive nucleotide sequences and differences in the karyotype structure C-banding pattern were used to confirm the absence of the M genome in *Ae*. *neglecta* and *Ae*. *columnaris* (Badaeva et al., 2018), which in this study was verified with thousands of loci. Therefore, we suggest research communities for the consistent use of genomes symbol for *Ae. columnaris* (UX) and *Ae*. *neglecta* (UX or UXN). Further cytological and genomic evaluation of the X genome is certainly required.

### *Aegilops* Genetic Diversity

The ploidy level and the mode of fertilization appeared as major determinants of *Aegilops* accessions diversity (Table 1). We did not observe the direct impact of population size on Nei’s diversity index (Nei, 1987) at any ploidy levels (Table 1). For instances, the diploid *Ae*. *sharonensis* (9 accessions) had a higher diversity index (0.019) than *Ae*. *umbellulata* (58 accessions); the tetraploid *Ae*. *ventricosa* (17 accessions) had a higher diversity index than another tetraploid *Ae*. *triuncialis* (199 accessions) (Table 1). As we observed the *Ae*. *speltoides* as the diploid species with the greatest diversity, we also observed relatively higher diversity indices of the S genome polyploids such as *Ae*. *kotschyi*, *Ae*. *peregrina* and *Ae*. *vavilovii* (Table 1). Overall, most of the *Aegilops* species had a wider and more variable diversity and had greater potential to utilize in wheat breeding. Therefore, serious efforts must be taken for the *in-situ* conservation of these germplasms and for enhancing *ex-situ Aegilops* germplasm collections. Kilian et al. (2011) also put forth the urgency of protecting these *Aegilops* germplasms and described the importance of understanding *Aegilops* genetic diversity, *Aegilops*-*Triticum* molecular biological relationships and identifying and preserving suitable *Aegilops* alleles for wheat breeding.

### *Aegilops* and Wheat Genomes

This is perhaps the first report showing genomic relationships between all *Aegilops* genomes and wheat sub-genomes based on high-throughput sequence-based markers and robust phylogeny of these wild wheat. As in some earlier reports, we observed that most of the *Aegilops* genomes (U, M, N, C) were genetically closer to the wheat D subgenome (Supplemental Figures S9 – S15) except *Ae*. *speltoides* (Figure 8). Several studies have reported that the B genome donor speciation event occurred earlier than the *Ae*. *tauschii* speciation (D-genome lineage) and therefore, there are stronger evolutionary relationships of the U, M, N and C diploid *Aegilops* within the D-genome lineage (Said et al., 2021; Tanaka et al., 2020; Glémin et al., 2019).

Still in the study, we observed unique relationships between some of the genomes of the *Aegilops*-*Triticum* complex which was not clearly described in earlier studies. One of the most important observations we made here is that four *Sitopsis* species have a relationship with both B and D subgenomes of wheat. The phylogenetic tree and the sequence read as well as loci-mapped statistics showed these relations (Supplemental Figures S8-S10). Nevertheless, other recent reports also considered these four members of the *Sitopsis* as (D lineage) and are closer to the wheat D subgenome (Li et al., 2022; Li, 2011; Avni et al., 2022).

### *Ae*. *mutica*, wheat genomes and homoploid hybridization

In this study, we observed unique genetic characteristics of *Ae*. *mutica* as it was phylogenetically closer to the *Ae*. *speltoides* (Figures 3-5 and Supplemental Figure S8), however, it showed genetic similarity with the wheat D subgenome (Supplemental Figure S15). Nevertheless, similar observations were also reported in recent studies. Li et al. (2022) reported lower genetic similarities between *Ae*. *mutica* and wheat B subgenome computed as the genetic relatedness. Grewal et al. (2022) reported a similar relation between *Ae*. *mutica* and wheat subgenomes as we reported here; where they found the highest *Ae*. *mutica* loci mapped on the D subgenome, rather than A and B subgenomes (Supplemental Figure S15). Therefore, the genetic similarities and phylogenetic relationship between the *Ae*. *mutica* and the *Aegilops*-*Triticum* complex are exclusive and should be studied in a larger population and high-depth sequencing. Also, these analyses indicate that *Ae*. *mutica* genome could have evolved itself or played a role in the evolution of polyploid genomes after its divergence from *Ae*. *speltoides*. Some recent studies also argued that *Ae*. *mutica* and the D lineage have homoploid hybridization followed by introgression (Bernhardt et al., 2020; Li et al., 2022). Bernhardt et al. (2020) reported that most of the members of the genus *Aegilops*, except *Ae*. *speltoides*, have evolved through ancient primordial hybrid speciation events between the ancestral *Triticum* and *Ae*. *mutica*. Very earlier studies also indicated the higher homology between *Ae*. *mutica* and the wheat D subgenome (Jones and Majisu, 1968).

### Utilizing *Aegilops* Novel Alleles in High-Throughput Genotyping Era

This study lays a solid foundation for the future utilization of *Aegilops* germplasm in the gene bank. With the development of introgression populations for breeding new genomics tools can enable rapid selection and advancement of novel alleles (Adhikari et al., 2022b). Likewise, association genomics approaches can be leveraged to identify novel alleles directly within the wild germplasm collections (Gaurav et al., 2022). In this context the importance of these highly diverse genetic resources is further enhanced. Finding trait-related alleles through genome-wide association studies (GWAS), generating reference assemblies and resequencing diverse panels are some future steps to utilize these valuable *Aegilops* genetic resources in wheat enhancement.

## Conclusions

In this study, we further uncovered the genomic and genetic relationships of all *Aegilops* species together and showed that the GBS approach can be used efficiently to curate the gene bank accessions and investigate the genetic diversity and population structure of the entire *Aegilops* collection. Most likely this is the first genomic analysis of a nearly complete set of the genus *Aegilops* with 23 species, where we dissected a larger population (1041) using over 45K SNPs and constructed a robust phylogenetic tree and the PCA clusters. The population grouping and structuring of this valuable wild wheat species mostly harmonized with the traditional nomenclatures at the species level. However, based on these high-throughput genome-wide markers, we also confirmed the genome symbols of two tetraploid species that were under controversies in the literature. Here we showed that each *Aegilops* subgenome and wheat subgenomes have unique relations at the genomic level, and require more investigation, especially *Ae*. *mutica* showed unique characteristics as appeared as the sister group of *Ae*. *speltoides* but more sequences and variants of this out-crossing species were mapped on wheat subgenome D. The genetic and evolutionary relations among *Aegilops* and with wheat will be clearer when we will have more genomic resources such as genomic assemblies and resequencing data for each of these *Aegilops* specie. Here we showed the relative diversities of all 23 species together for the first time. The substantial genetic diversity and its relative extent in each of the *Aegilops* species provided an opportunity to select species and germplasms as sources of novel alleles for wheat breeding and improvement.

## Materials and Methods

### Plant Resources

This study primarily included 1041 accessions of the *Aegilops* species preserved and maintained in the WGRC gene bank (Supplemental Table S1 and Figure 1). The accessions were originally collected from various sources and sites including the Middle East, Anatolia, East Asia, and Northern Africa (Figure 1 and Table S1). Accessions comprise 22 different *Aegilops* species under five sections (*Aegilops*, *Comopyrum*, *Cylindricum*, *Sitopsis* and *Vertebrata*) (van Slageren, 1994) and *Ae*. *mutica,* which is synonymously known as *Amblopyrum muticum*. For gene bank curation and most part of the population analysis, only those *Ae*. *tauschii* accessions that were not in the previous gene bank curation experiment (Singh et al., 2019a) were used. We also used CIMMYT wheat lines and already curated *Ae*. *tauschii* lines (Table S1) for genotyping together with the diploid *Aegilops* to dissect the genetic relationships among wheat and *Aegilops* genomes.

Most of these species are self-pollinated and were primarily maintained by single seed descent, with exceptions described below. *Ae*. *speltoides* and *Ae*. *mutica* which are partially outcrossing and were maintained through sib-mating multiple plants. Specifically, *Ae*. *mutica* accessions consisted of 54 samples from five out-crossing plants bagged together.

### Genotyping and Markers Identification

The DNA extraction, GBS library preparation and sequencing were performed as we described in our earlier studies (Adhikari et al., 2022a) using two enzyme-based GBS (Poland et al., 2012). Only a single plant per accession was sequenced for all species except *Ae*. *mutica* where we sequenced 54 individuals obtained from randomly crossing five plants because the species is cross-pollinating and it has a low germination rate.

For the *de novo* SNP calling, reads were demultiplexed using sabre (https://github.com/najoshi/sabre) and adapters were trimmed using fastp (Chen et al., 2018). The variants were called using the available reference assemblies of diploid *Aegilops* and wheat, and using mock references generated as described (Melo et al., 2016; Adhikari et al., 2018). For mock references, the raw GBS reads of selected accessions with higher sequence data were used as the reference source. We also ensured that the mock reference represents the sequences of relevant *Aegilops* species or the genomes [C, D, M, N, S, U, T] for the population to be genotyped. The *de novo* variants were called using BCFtools (Li, 2011) and used for initial gene bank curation and population clustering of the whole collection. Then the *de novo* variants were also called for some species independently depending on the objectives of the specific analysis (Table S3). For some species in polyploid lineages, we called variants on a diploid ancestor and later the same variants were called in the polyploids using BCFtools (Li, 2011). After calling variants, unless otherwise stated, we filtered loci to keep any variants passing these conditions; minor allele frequency (MAF) > 0.01, missing < 30%, and heterozygous < 10%.

The TASSEL5 GBSv2 pipeline was used for reference-based SNP calling (Glaubitz et al., 2014). For this method, *Ae*. *tauschii* reference genome Aet v5.0 (Wang et al., 2021) or *Ae*. *sharonensis* (Yu et al., 2022) *Ae*. *speltoides* (Avni et al., 2022), *Ae*. *searsii* and *Ae*. *bicornis* ((Li et al., 2022) genomes were used. We also called variants in all these diploids species to the wheat reference using the ‘Chinese Spring’ wheat reference (IWGSC CS RefSeq v2.1) (Zhu et al., 2021) to observe the relationship between *Aegilops* and wheat.

### Gene Bank Curation

In the first step, the germplasm curation identified misclassified accessions and corrected the taxonomy of these accessions in the database (Singh et al., 2019a). We identified misclassified accessions by constructing a phylogenetic cluster colored with the recorded species. These were further verified using PCA clustering followed by a visual assessment of seeds and spikes. The misclassified accessions were identified and confirmed with multiple genotyping sets viz. entire collection, species alone and same genome accessions together.

In the second step, the genetically identical accessions were determined using allele matching (Singh et al., 2019a; Adhikari et al., 2022a). However, this assessment was done only for the accessions of the species whose reference genome is available e.g., *Ae*. *tauschii* and the *Sitopsis* section *Aegilops*. The allele matching (> 99% identity-by-state) was used as a threshold to confirm genetically identical accessions. Allele matching used homozygous and non-missing sites between two given accessions, and the raw markers were filtered using MAF > 0.01, missing < 50%, and heterozygous < 20% parameters before allele matching.

### Genetic Clustering, Population Analysis and Diversity

The genotyping matrices were analyzed for the genetic distances among the *Aegilops* populations, which were then used for exploring the population structure and ancestry. For phylogenetic clustering, the genetic distance was computed using the ‘dist’ function in R (R language 2020) and the R packages *ape* (Paradis and Schliep, 2019) and *pyclust* (Chen, 2011) were then used to generate unrooted neighbor-joining (NJ) tree with the default parameters (Singh et al., 2019b; Adhikari et al., 2022a).

The genetic relationships among the *Aegilops* accessions were further examined via principal component analysis (PCA), which was performed in two steps. The A matrix was derived from A.mat() function within the R package rrBLUP (Endelman, 2011) and the eigenvalues and eigenvectors were derived using the ‘e’ function as in (Adhikari et al., 2022a). Further, the population structure of the *Sitopsis* group of *Aegilops* was also performed with the reference-based genotyping profile using fastStructure software (Raj et al., 2014) as explained (Adhikari et al., 2022a). We computed Nei’s diversity index (Nei, 1987) and total segregating loci for each of the *Aegilops* species to assess the relative diversity of the species.

### Ae. columnaris and Ae. neglecta Genome Symbols

We investigated the traditional genome symbols of *Ae*. *columnaris* (UM) and *Ae*. *neglecta* (UM, UMN) for the presence/absence of the M genome. There are recent cytology-based findings that have questioned the traditional genome symbols of these species (Badaeva et al., 2018). To test this, we computed the sequence read mapping and segregating loci on the M and U mock reference genomes for the *Ae*. *columnaris* and *Ae*. *neglecta* accessions as well as two other tetraploids (*Ae*. *nelglecta* and *biuncialis*) whose genomic compositions are unequivocally recognized as MU or UM. The *de novo* variants were first identified for the diploid M genome (*Ae*. *comosa*) and U genome (*Ae*. *umbellulata*) populations separately, and then the same variants were called on these four tetraploid species. We also constructed the phylogenetic clustering among *Ae*. *columnaris*, *Ae*. *neglecta*, *Ae*. *geniculata*, *Ae*. *biuncialis* and a tetraploid that shares only the U genome (*Ae*. *triuncialis*) to see their relative positions in the tree (Supplemental Figure S3).

### *Aegilops* Genome Relation to Wheat Genome

We mapped diploid *Aegilops* GBS reads to the wheat genome (CS.Ref.v1) (Appels et al., 2018) and computed sequence read mapping coverage. The reads mapped per Mb wheat subgenome and the total variants mapped for each wheat subgenomes (A, B, D) were recorded. We did not further evaluate *Ae*. *tauschii* whose close genetic relationship as the wheat D subgenome donor has been clearly established. We also generated an unrooted neighbor-joining (NJ) phylogenetic tree among diploid *Aegilops* and wheat using the variants called on wheat B and D reference subgenomes independently.

### Accession Numbers

The Raw GBS data, the fastq files, are available in the Sequence Read Archive (SRA) of National Center for Biotechnology Information (NCBI) under the BioProject accession PRJNA985892. The key file and necessary SNP matrices and the R script files (.rmd) are provided in the dryad public repository which are available with the unique DOI: 10.5061/dryad.mgqnk994n.

All data are available in the manuscript or the supplementary files and at the Dryad digital repositories https://datadryad.org/stash/dataset/doi:10.5061/dryad.mgqnk994n

## Supplemental Data

**Supplemental Figure S1**. The GBS SNP based unrooted neighbor-joining (NJ) tree separating tetraploid and hexaploid accessions of *Ae*. *neglecta* (blue clade) and the chromosome counts of two representative individuals from each 4X and 6X sub-clade of the *Ae*. *neglecta*.

**Supplemental Figure S2**. Principal component analysis (PCA) plot showing two forms of *Ae*. speltoides; var. *speltoides* and *ligustica*.

**Supplemental Figure S3**. An unrooted neighbor-joining (NJ) tree separating some tetraploid *Aegilops* accessions containing two species whose genome formula is controversial, the *Ae*. *neglecta* and *Ae*. *columnaris*.

**Supplemental Figure S4**. The bar chart showing the overall sequence read alignment of four tetraploid *Aegilops* species; *Ae. biuncialis, Ae*. *geniculata, Ae*. *columnaris* and *Ae*. *neglecta* when aligned on M and U genome *de novo* mock reference.

**Supplemental Figure S5**. An unrooted neighbor-joining (NJ) tree of *Ae*. *juvenalis*, *Ae*. *crassa and Ae*. *vavilovii.* The tree branches were colored based on the accession’s taxon.

**Supplemental Figure S6**. Minor allele frequency (MAF) distribution within the loci for the entire *Aegilops* collection.

**Supplemental Figure S7**. Distribution of minor alleles frequency (MAF) for segregating variants in *Ae*. *speltoides*.

**Supplemental Figure S8**. An unrooted neighbor-joining (NJ) tree constructed using the genotyping information generated by using wheat B genome as a reference (left); and the unrooted NJ tree constructed using genotyping profile generated using the wheat D genome as a reference (right).

**Supplemental Figure S9**. Bar charts showing genomic relations between the *Sitopsis* section *Aegilops* (except *Ae*. *speltoides*) and the wheat.

**Supplemental Figure S10**. Bar charts showing genomic relations between the *Sitopsis* section *Aegilops* (except *Ae*. *speltoides*) and the wheat.

**Supplemental Figure S11**. Bar chart showing genomic relation between U genome diploid *Ae*. *umbellulata* and wheat.

**Supplemental Figure S12**. Bar chart showing genomic relation between N genome diploid *Ae*. *uniaristata* and wheat.

**Supplemental Figure S13**. Bar chart showing genomic relation between M genome diploid *Ae*. *comosa* and wheat.

**Supplemental Figure S14**. Bar chart showing genomic relation between C genome diploid *Ae*. *markgraffii* and wheat.

**Supplemental Figure S15**. Bar charts showing genomic relations between *Ae*. *mutica* and wheat.

**Supplemental Table S1**. List of *Aegilops* germplasms in the WGRC gene bank collection with the taxa and origins of the accessions [separate excel file].

**Supplemental Table S2**. Misclassified and genetically identical (redundant) *Aegilops* accessions [separate excel file].

**Supplemental Table S3**. Different SNP matrices, population genotyped, the reference sequence used and the application which used the SNP matrix.

**Supplemental Table S4.** *Sitopsis* section *Aegilops* and *Ae*. *mutica* pairwise F_ST_ values.

**Supplemental Table S5**. Total segregating loci in UM and UX genome species when called variants on the M genome and U genome mock references independently.

## Acknowledgements

We would like to acknowledge Kansas high-performance computing cluster ‘beocat’ for data storage and the Linux environment for data analysis. We are thankful to everyone who contributed to WGRC gene bank *Aegilops* collection.

